# Neural network and random forest models in protein function prediction

**DOI:** 10.1101/690271

**Authors:** Kai Hakala, Suwisa Kaewphan, Jari Björne, Farrokh Mehryary, Hans Moen, Martti Tolvanen, Tapio Salakoski, Filip Ginter

**Author notes:** These authors contributed equally to this work (Kai Hakala, Suwisa Kaewphan, Jari Björne). Department of Future Technologies, University of Turku, Turku, Finland.

## Abstract

Over the past decade, the demand for automated protein function prediction has increased due to the volume of newly sequenced proteins. In this paper, we address the function prediction task by developing an ensemble system automatically assigning Gene Ontology (GO) terms to the given input protein sequence.

We develop an ensemble system which combines the GO predictions made by random forest (RF) and neural network (NN) classifiers. Both RF and NN models rely on features derived from BLAST sequence alignments, taxonomy and protein signature analysis tools. In addition, we report on experiments with a NN model that directly analyzes the amino acid sequence as its sole input, using a convolutional layer. The Swiss-Prot database is used as the training and evaluation data.

In the CAFA3 evaluation, which relies on experimental verification of the functional predictions, our submitted ensemble model demonstrates competitive performance ranking among top-10 best-performing systems out of over 100 submitted systems. In this paper, we evaluate and further improve the CAFA3-submitted system. Our machine learning models together with the data pre-processing and feature generation tools are publicly available as an open source software at https://github.com/TurkuNLP/CAFA3

**Author summary:** Understanding the role and function of proteins in biological processes is fundamental for new biological discoveries. Whereas modern sequencing methods have led to a rapid growth of protein databases, the function of these sequences is often unknown and expensive to determine experimentally. This has spurred a lot of interest in predictive modelling of protein functions.

We develop a machine learning system for annotating protein sequences with functional definitions selected from a vast set of predefined functions. The approach is based on a combination of neural network and random forest classifiers with features covering structural and taxonomic properties and sequence similarity. The system is thoroughly evaluated on a large set of manually curated functional annotations and shows competitive performance in comparison to other suggested approaches. We also analyze the predictions for different functional annotation and taxonomy categories and measure the importance of different features for the task. This analysis reveals that the system is particularly efficient for bacterial protein sequences.

## Introduction

Proteins play a pivotal role in many processes of living organisms, including, but not limited to, signal transduction, transmembrane transport and structural support. Determining protein functions experimentally is an expensive and labor-intensive undertaking. With the increasing number of sequences produced by high throughput sequencing methods, there is an urgent need for computational methods to assist in protein function annotation. Over the past decade, a research community focusing on automated function prediction (AFP) has formed, resulting in a number of AFP systems and the regular Critical Assessment of Functional Annotation (CAFA) challenge.

CAFA is a shared task organized by the AFP-Special Interest Group, aiming to establish a common platform and evaluation methods for measuring the performance of automated systems for the AFP task [1, 2]. In CAFA, each participating research group is asked to predict the functions for a large set of proteins with more than 100,000 individual sequences, under a tight time limit. Subsequently, the organizers contract laboratories to experimentally verify functional annotations for a subset of the sequences, usually within half a year after the predictions were submitted. The resulting verified annotations constitute a new test set, against which the predictions made by the participants can be evaluated.

In CAFA, the protein functions are assigned from the controlled vocabulary of terms defined in Gene Ontology (GO), i.e. the task is to annotate the given protein sequences with the relevant GO terms that describe their function. In addition to molecular functions (MF), GO includes terms related to the cellular components (CC) in which the proteins are active, and the biological processes (BP) such as pathways in which the proteins participate [3, 4]. All together, over 40,000 terms, organized in a hierarchy, exist in the current version of GO. This translates into a multi-class and multi-label classification task, where each protein sequence can be annotated with multiple terms from this large vocabulary, with strong statistical dependencies between the terms. Further, the distribution of GO terms is highly skewed in several distinct ways, as demonstrated in Swiss-Prot, a subset of the UniProt protein database [5] manually curated with GO terms. While for instance the human proteins are densely annotated with 20 GO terms on average, full 36% of GO terms do not have a single annotated example, and 18% have only one. This means that a proportionally small number of unique GO terms account for a large proportion of the annotations. All these factors combined make AFP a very challenging task from the machine learning perspective.

The first two CAFA challenges have seen a variety of approaches applied to the problem [1, 2]. Among the top performing systems, the most common approach was the annotation transfer by homology, combined with a statistical or machine-learned scoring function. Cozzetto et al. [6] use a scoring function to combine and rank the predictions from various biological data analyses, including PSI-BLAST [7], profile-profile comparison, text mining, sequence features, protein-protein interactions and high-throughput data. This approach was ranked as the best performing system in CAFA1, which evaluated only the *molecular function* and *biological process* GO ontologies. The PANNZER method [8] uses weighted k-nearest neighbor to predict protein functions from the weighted sequence similarity scores using BLAST [9] search, and the taxonomic distance of the organisms originating the sequences. Argot2 [10] uses the semantic similarity of weighted ontology terms found through the BLAST sequence similarity search and the HMMER [11] tools. In CAFA2, GO-FDR [12], one of the best performing systems on all three GO ontologies, calculates the probability of a protein being associated with a target GO term, using predictions from the PSI-BLAST tool.

Recently, You et al. [13] suggested an approach based on an ensemble of logistic regression models which, resulted in the best overall performance among the participating teams in the CAFA3 challenge. For each GO term, a set of three logistic regression models are independently trained based on structural information from InterPro [14], biophysical attributes from ProFET [15] and amino acid n-gram features. Sequence alignment and GO annotation frequencies are used as additional features. All this information is aggregated by a separate machine learning (ML) model in a learning-to-rank setting.

Deep neural network architectures have been successfully applied to bioinformatics problems, such as fold recognition, functional classification and protein design [16, 17]. For the protein function prediction task, several architectures have been developed, aiming to replace hand-crafted features with ones directly extracted from the sequences. DeepGO [18] uses a deep convolutional neural network (CNN) architecture composed of 10 hidden layers, with inputs formed from amino acid tri-grams and the embedding (continuous vector representation) of the protein induced using its neighborhood in the STRING [19] database graph. ProLanGO [20] uses a neural machine translation system based on a recurrent neural network to “translate” proteins into the corresponding GO terms. Here the protein sequences are first converted to non-overlapping amino acid k-mers of length 3 to 5, forming the input sequence for the model. Rui et al. [21] use multi-task deep neural networks with features directly derived from the amino acid sequences as well as external analysis tools, focusing on human proteins only. In general, these deep neural network models have demonstrated a competitive performance, on par with other machine learning approaches with less feature engineering effort.

Random forests, an ensemble of decision trees, have been used in a wide array of prediction tasks, including post-translational modification site prediction [22], fold recognition [23], SCOP structural classification [24], protein-protein interaction site prediction [25], and enzyme function classification [26]. Most models derive protein-related features, such as hydrophobicity, secondary structure, and amino acid composition, both by directly extracting the features from the sequence and by relying on established sequence analysis tools. Random forests have been a popular machine learning method in bioinformatics due to its simplicity in terms of modeling and interpretability of the results through feature importance analysis.

In this paper, we introduce our system based on neural network and random forest classifiers, both relying on a rich set of features and, individually, achieving competitive performance. Further, through an evaluation on our internal test dataset, we show that these individual approaches strengthen each others performance, resulting in an ensemble system outperforming the two individual classifiers. For comparison, we also experiment with a neural network relying purely on the sequence itself, in order to evaluate the feature generation ability of the neural networks. Most importantly, this neural network does not have access to the manually curated annotations of homologous proteins, a primary source of features for all systems performing well on the AFP task, nor to the biologically motivated features produced by established sequence analysis tools.

The official CAFA3 evaluation places our ensemble system in top-10 overall out of 68 participating teams and 144 submitted systems, with particularly strong performance on molecular function and cellular component categories of prokaryotic proteins, where the system placed 3rd and 2nd [27].

## Materials and methods

In this section we describe the protein datasets, sequence analyses used for generating features and the details of the neural network and random forest classifiers and the ensemble systems.

### Protein dataset

As the *training data*, we selected only those Swiss-Prot proteins, whose function annotation is based on a reliable experimental evidence, which we define as evidence codes `EXP, IDA, IPI, IMP, IGI, IEP`, the author statement code `TAS`, or the curatorial statement evidence code `IC` being assigned to the annotation. This restriction aims to discard annotations stemming from high-throughput and other noise-prone experimental techniques. The training data consists of 387,416 individual annotations of 67,118 proteins, from the total of 738,431 annotations of 112,279 proteins in Swiss-Prot that have other than purely computationally predicted annotation.

In addition to the exact GO terms manually assigned to a particular protein, we enriched the annotations by additionally assigning the ancestral terms from the ontology to the proteins, increasing the number of individual annotations to 3,955,953. As a result, the proteins which are on average annotated with 6 terms, now become annotated with 60 terms.

### Sequence analysis and protein features

In this section, we summarize the sequence-based features as used by the various classifiers throughout the paper.

#### BLAST features

We use the Protein-Protein BLAST (BLASTP) program from a locally installed NCBI-BLAST+ version 2.5.0 [28] to search for similar proteins from the full Swiss-Prot database. The E-value of 0.001 is used as the inclusion threshold. We query with both BLOSUM45 and BLOSUM62 [29] scoring matrices, resulting in two distinct sets of similar proteins for each query sequence. These will be referred to as *blast45* and *blast62* hereafter. We use the default gap scores for each of the scoring matrices.

We also use DELTA-BLAST (Domain Enhanced Lookup Time Accelerated) for searching distantly related protein sequences [30]. DELTA-BLAST increases the sensitivity of BLAST-based sequence similarity search by constructing position-specific score matrices from the conserved domain database (CDD) [31]. BLOSUM62 is used as the scoring matrix with 0.001 as the cut-off E-value.

All BLAST data was converted to feature vectors with the following approach: The Uniprot ID of the matched protein was used as the feature name and the HSP (High-scoring Segment Pair) score as its value.

#### InterproScan

InterProScan [32] is a software package predicting structural motifs, functional domains, signatures, protein families and other features relevant to protein function analysis, based on the InterPro database. We use locally installed InterProScan 5 to predict InterPro profiles and, subsequently, GO terms for those cases where a mapping between an InterPro profile and GO is established in the database. These mappings are however neither up-to-date nor complete. To increase the coverage of the mappings, we trace the patterns and signatures to InterPro’s upstream databases, and recover the GO terms from there. Features are generated from the matching GO terms (or a special feature is produced, signalling the absence of such mapping) such that scores given by InterProScan are converted to numerical features while the *GO* terms and profile *accession* identifiers are converted to binary features.

#### Taxonomy features

The NCBI Taxonomy is a manually curated database of names and taxonomic lineages for organisms within the scope of the International Nucleotide Sequence Database Collaboration (INSDC) [33]. A binary feature is generated for each node in the NCBI Taxonomy and subsequently assigned to each protein based on its organism of origin.

#### Sequence features

We use several additional tools to analyze the protein sequence and provide features potentially relevant to its function.

For nuclear localization, we use NucPred [34] to predict whether a protein enters the nuclear compartment. This analysis applies eukaryotic sequences only, as prokaryotes do not have a nucleus. For post-translational modifications, we use NetAcet [35] to predict proteins which are acetylated by N-acetyltransferase A (NatA). For GPI-anchored proteins, we use PredGPI [36] which is based on a support vector machine and a Hidden Markov Model predicting the anchoring signal and the most probable omega-site. The predictions of these three tools are encoded as numeric features.

#### Amino acid index

The amino acid index is a set of numerical values characterizing the physicochemical and biochemical properties of each of the 20 amino acids. It has been used for numerous structure-function prediction tasks, e.g. human protein sub-cellular localization [37]. In this work, we obtain 544 amino acid indices from the Amino Acid Index database [38] and use their numerical values as features in our experiments with the sequence-only convolutional neural network (See Section “Feedforward neural network classifier”).

### Experimental setting

In all experiments the input data for each protein was converted into numerical feature vectors and the GO terms into binary label vectors. We subsequently randomly divide the training data into three parts: The *training* set used for training the classifiers, the *validation* set used for hyperparameter optimization and the *test* set which is used for the final performance estimation. The validation set is also used during system development and for experiments, including feature selection, in order to avoid overfitting the test set. The training, validation and test subsets contain 60%, 20% and 20% of the whole data.

Different feature groups were tested in combination to select the ones that gave the best classification performance. Finally, for computational reasons, the classifiers are trained to predict the 5,000 most common terms, a subset of GO which covers over 94% of all GO annotations in Swiss-Prot. Note that in testing, the classifiers are naturally evaluated on the full set of GO terms, thereby the 5,000 term subsetting choice does not artificially overestimate the performance.

We evaluate the performance of the systems using the F-score, unlike in the official CAFA evaluation, where the maximal F-score, calculated from precision-recall curves, is used as the primary metric. Thus, our internal evaluation is more strict and acts as a lower bound for the maximal F-score.

### Feedforward neural network classifier

As the first classifier in the ensemble, we train a standard feedforward neural network receiving a vector of the features described in Section “Sequence analysis and protein features”. The feature values are scaled by dividing them by the maximum absolute value observed in the *training* set for the given feature, before they are utilized in the model. This scaling has favorable computational implications, as zero values are preserved as such, and feature matrices can be stored in the sparse format. To further reduce the computational cost, we reduce the number of features by removing those with variance below 0.0001 after scaling. The input features are passed to a fully connected layer with dimensionality of 300 and hyperbolic tangent activation function before the output layer, which is a standard dense layer of dimensionality 5,000, corresponding to the number of unique output GO terms. Since the learning task is multi-label, i.e. the classifier can predict several classes for each instance, the output layer is trained using the binary cross entropy objective.

The network is regularized by applying a dropout with rate 0.5 to the input features [39]. The training is stopped early, once the *validation* set F-score is no longer improving.

The substantial imbalance towards the negative class in the multi-label mode of training results in a high precision and low recall model. To mitigate this, the recall is boosted by penalizing false negative predictions more than false positive ones. The magnitude of the penalty is a hyperparameter selected to optimize the performance on the *validation* dataset.

### Random forest method

Random forests [40] are a prediction algorithm based on an ensemble of decision trees. Each decision tree in the ensemble is built based on a different random subset of the input features, and the final prediction of the ensemble is the majority vote among the trees. Random forests are particularly suitable for the current task as they have good classification performance and support multi-label classification on thousands of labels. The classifier was implemented using scikit-learn library version 0.18.1 [41].

### Convolutional neural network

While the previous two methods can be seen as traditional classifiers relying on a carefully selected set of features, the third method, convolutional neural networks, will depart from this paradigm in not being presented with any complex features. Rather, the neural network is presented with the sequence itself as its sole input.

Convolutional neural networks (CNNs) have been successfully applied to sequential and multidimensional prediction tasks, especially in computer vision and text classification [42, 43]. In biological sequence analysis, they have been applied specifically to protein secondary structure prediction [44]. The strength of CNNs arises from the possibility of detecting fuzzy patterns locally from the input data, e.g. in our case we expect the convolutional kernels learn to detect amino acid n-grams relevant to a certain protein function. Many suggested systems for AFP rely on structural information such as the presence of a certain domain or motif. However, this type of information is mostly gathered from other tools such as InterProScan, or by simply looking at the amino acid n-grams present in a given sequence, resulting in extremely sparse feature representations [18]. To overcome this issue, our neural network learns a latent feature vector (embedding) for each unique amino acid, and a protein is represented as a sequence of these embeddings. This gives the model an opportunity to measure whether two different amino acids tend to have a similar role in the sequence, leading to similar embeddings. To ease the task of learning these embeddings, we attempted to initialize the weights with the Amino Acid Index properties, but this did not have a significant influence on the model performance.

We use convolution window sizes of 3, 9, 27 and 81 amino acids and learn 50 kernels for each window size. The shortest window size of 3 is a common length used in amino acid n-gram features, whereas size 9 approximates the length of local segments analyzed in protein secondary structure prediction [44]. The larger window sizes of 27 and 81 should in turn be able to detect motifs and shorter domains [45].

The convolutional kernel activations are subsequently pooled by taking the maximum activations of each kernel across all positions in the sequence (max pooling). This procedure removes the location information of the detected patterns, producing a fixed-length, position-invariant model suitable as an input to a classification layer. A prior study shows that trying to preserve the location information with local max pooling does not provide any benefits in DNA-protein binding prediction [46], hence we have not experimented with other possible settings.

These maximum kernel activations then form an input of a fully connected output layer, producing the final predictions. The dimensionality of the output layer is 5,000, corresponding to the number of predicted GO terms.

Proteins longer than 2500 amino acids are truncated, i.e. we analyze only the first 2500 amino acids of each sequence. This truncation affects only 1% of the training sequences.

### Homology transfer

In our previous work on the CAFA2 challenge [2], we observed that oftentimes proteins received no prediction from the classifiers, despite there being homologous proteins with existing annotation. Moreover, our classifiers are trained with only the top-5000 most common annotated terms, leaving less frequent terms unattended. To address the issue, we use the following simple fallback homology-based transfer approach, inferring the functions from homologous proteins. For each protein, we extract the GO terms associated with homologous sequences from the *blast62* alignment, i.e. all sequences identified as similar at the BLAST E-value cut-off of 0.001. We subsequently rank the ontology terms by the number of associated homologous sequences, and the top 5 terms which are supported by at least 2 sequences form the prediction of the Homology Transfer fallback method.

### Ensemble

As the final, combined output of the abovementioned methods, we take the union of their predictions. We evaluate all the model combinations and use the subset of models that leads to the best performance measured in terms of F-score as the final system. For completeness, we will also report on results obtained using the intersection of the predictions, a distinctly high precision, low recall model. The whole system architecture is visualized in Fig 1.

**Fig 1.**
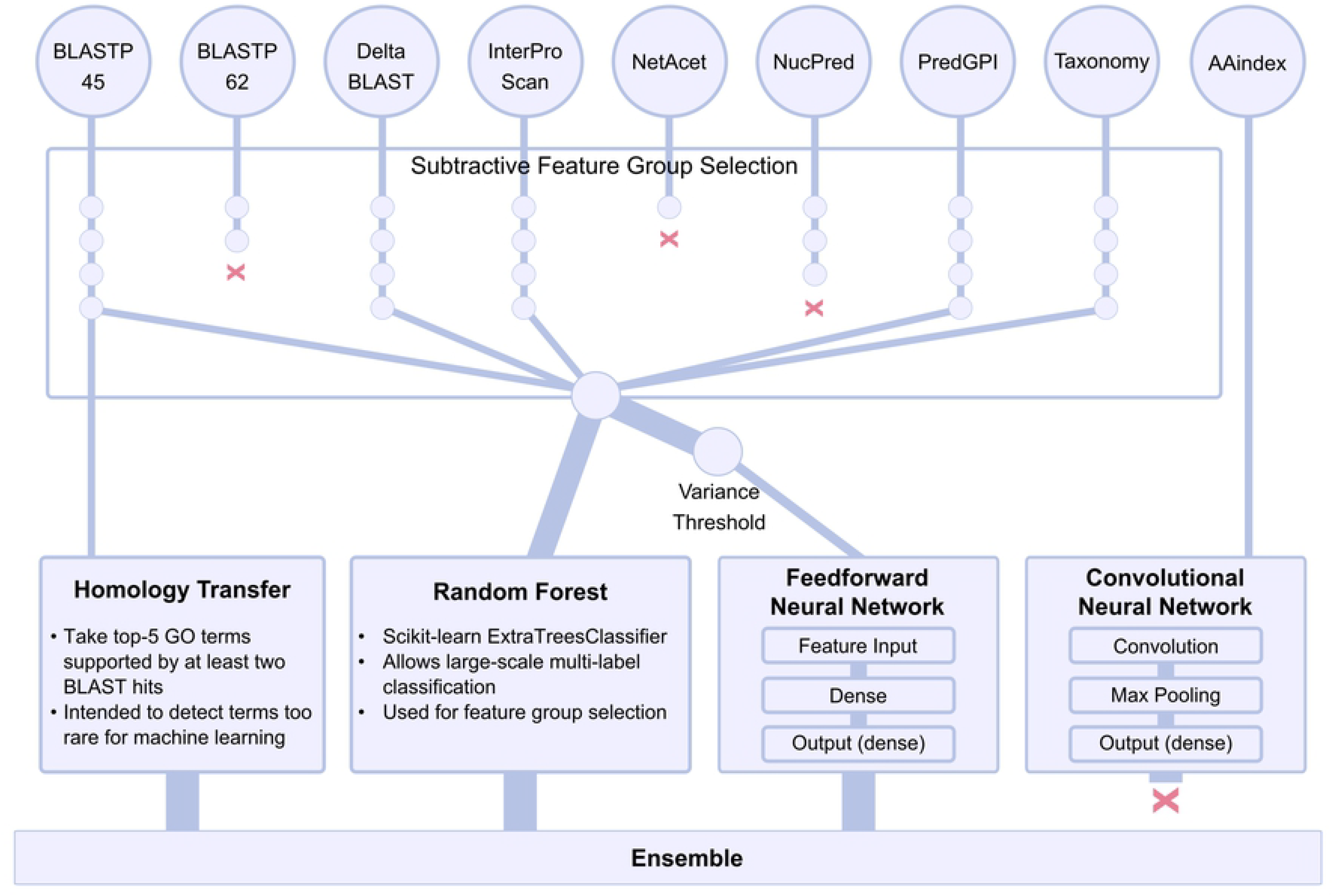
System architecture. The architecture of our ensemble system, based on three approaches, homology transfer, random forest and neural network. Features from Blast62, NetAcet and NucPred analyses are removed from the final feature sets. The final set of features are used in both random forest and machine learning systems. The final ensemble system is a union (boolean *OR* combination) of the predictions from the three approaches.

## Results

In this section, we evaluate the methods from several distinct angles as well as report on the ranking of the ensemble system in the CAFA3 challenge.

### Feature group selection

Feature selection is a process of selecting a subset of relevant features, or removing irrelevant features, in order to simplify the system, reduce the training time and improve the generalization of the models [47]. We performed feature selection by repeatedly removing one feature group at a time, so as to increase the overall performance on the development dataset, stopping when the performance of the classifier no longer increased (see S1 Table for detailed results). The optimal performance using the random forest classifier on the development data was achieved by removing the *blast62*, *netacet* and *nucpred* feature groups. The same feature subset is used also for the neural model.

Among the remaining features, *taxonomy* contributes the most to the system performance, i.e. removing *taxonomy* features results in a substantial drop in F-score. For BLAST-based homology features, the fact that the *blast45* features outperformed the *blast62* features can be attributed to the BLOSUM45 matrix allowing more distant hits, thus increasing the recall compared to BLOSUM62. These results also emphasize that choosing the right scoring matrix can have an impact on the performance of the classifiers. In general, the selection of a BLAST scoring matrix depends on the proteins at hand and the downstream application.

### Method evaluation

We evaluate the performance of all systems against the test dataset using the precision/recall/F-score metric for both single and ensemble systems. The evaluation results on all of the 29,190 unique annotated GO terms are summarized in Table 1.

**Table 1.** The performance of all system combinations in this work.

| NN | RF | HT | mode | F | P | R |
| --- | --- | --- | --- | --- | --- | --- |
| CNN |  |  |  | 0.347 | 0.316 | 0.385 |
| FFN |  |  |  | **0.480** | 0.492 | 0.468 |
|  | RF |  |  | 0.424 | 0.609 | 0.326 |
|  |  | HT |  | 0.387 | 0.550 | 0.298 |
| FFN | RF |  | OR | **0.493** | 0.472 | 0.517 |
| FFN |  | HT | OR | 0.487 | 0.460 | 0.518 |
|  | RF | HT | OR | 0.471 | 0.527 | 0.426 |
| FFN | RF |  | AND | 0.398 | 0.707 | 0.277 |
| FFN |  | HT | AND | 0.363 | 0.675 | 0.248 |
|  | RF | HT | AND | 0.312 | 0.740 | 0.198 |
| FFN | RF | HT | OR | **0.493** | 0.445 | 0.553 |
| FFN | RF | HT | AND | 0.296 | 0.765 | 0.184 |

Table notes All ensemble combinations of the three methods: random forest classifier (RF), feedforward neural network (FFN), and the homology transfer (HT), using either intersection *AND* or union *OR* of the predictions. The performance is evaluated as the micro-averaged F-score (F), precision (P) and recall (R).

The performances of all the approaches, feedforward neural network (FFN), convolutional neural network (CNN), random forest classifier (RF) and homology transfer (HT) are shown in Table 1. The performance of all classifiers, except for CNN, surpass HT that is based solely on inferring known protein functions from similar sequences. FFN is the best performing method with an F-score of 0.480, followed by RF with 0.424.

The union of the predictions of the individual methods outperforms each method individually. The best result in terms of F-score is achieved by combining the predictions of FFN and RF. This demonstrates that the classifiers, even though provided with the same features, learn different aspects of the task. Also, as the RF classifier performs at the high precision - low recall point, it adds a small number of predictions, which are on average more likely correct than FFN, thereby benefiting the overall numerically stronger FFN method. Even though adding the predictions from HT improves F-score of both RF (+4.7pp) and FFN (+0.7pp) methods in isolation, it no longer improves the F-score of the RF-FFN ensemble. As expected, the intersection-based ensemble produces numerically inferior F-score as it drastically decreases the already low recall of the models. Nevertheless, this low recall is matched with a comparatively high precision, which could be a desirable property in some applications.

Finally, the CNN method has the lowest performance of the four methods in isolation, and also decreased the performance of all tested ensembles. These are therefore excluded from the Table 2 (see S2 Table for these results). Nevertheless, keeping in mind the minimal input information presented to the CNN classifier — the raw sequence itself — we find it very encouraging, even surprising that it can reach a performance which, in terms of F-score is roughly comparable to the HT method (CNN=0.347, HT=0.387) and a mere 15 percent points behind an ensemble of several strong methods with large feature sets. We have experimented with more complex CNN architectures, but neither increasing the kernel window, nor stacking several convolutional layers resulted in an improvement.

**Table 2.** The performance of the submitted systems to the CAFA3 challenge.

| NN | RF | HT | mode | F | P | R |
| --- | --- | --- | --- | --- | --- | --- |
| NN |  |  |  | 0.449 | 0.485 | 0.419 |
|  | RF | HT | OR | 0.471 | 0.527 | 0.426 |
| NN | RF | HT | OR | **0.483** | 0.442 | 0.532 |

### CAFA3

The ensemble system as submitted to the CAFA3 challenge differs from the methods above in not treating the FFN and CNN methods independently. Rather the outputs from the max-pooled CNN layer and the first fully connected layer of the FFN method are concatenated and serve as an input to a single output layer. In subsequent experiments we however found that an improvement of +3.1pp F-score can be achieved by removing the CNN component. The performance of the system submitted to CAFA3 is reported in Table 2. The detailed official results of CAFA3 are not available at the time of writing, therefore only our internal evaluation of the system is reported. Note that the values in Tables 1 and 2 are numerically comparable.

### Evaluation on model organisms and different ontologies

The ultimate goal for automated function prediction is to develop universal and reliable algorithms capable of predicting the function of any protein sequence. This is of course a very difficult task. As shown in all CAFA challenges [1, 2, 27] the accuracy of the predictions varies greatly among the organisms and different ontologies. To look beyond overall system performance, we next focus on the results of our methods on these two aspects: ontologies and organisms.

There are 10 organisms with over 1,000 manually annotated sequences, hereafter called *model organisms*. The list includes both domains of life, eukaryota (*Arabidopsis thaliana*, *Mus musculus*, *Rattus norvegicus*, *Homo sapiens*, *Drosophila melanogastor* and *Caenorhabditis elegans*) and prokaryota (*Schizosaccharomyces pombe* 972h-, *Saccharomyces cerevisiae* S288c, *Escherichia coli* K-12, *Mycobacterium tuberculosis* H37Rv). The number of annotated sequences from these 10 organisms, ranging from 1,500 for *M. tuberculosis* to 14,000 sequences for human, account for 83% of the whole protein dataset. We compare the performance of the classification methods with the homology transfer approach on the *model organisms* in the Swiss-Prot dataset.

On one hand, the improvement of the performance is only minor to moderate for multiple cellular organisms, ranging from 4pp to less than 15pp. As shown in Fig 2, proteins from *C. elegans* and mouse are more difficult to predict, as the systems add only +5pp on top of the HT F-score. On the other hand, the systems show higher performance improvement on bacterial sequences, increasing the F-score by 4–23pp, with the best overall improvement seen on *E. coli K-12* proteins. Despite having less annotated sequences, predicting functions of prokaryote proteins seems an easier task for machine learning systems. This is probably due to the fact that prokaryotes are simpler organisms with fewer functions and shallower GO ontologies.

**Fig 2.**
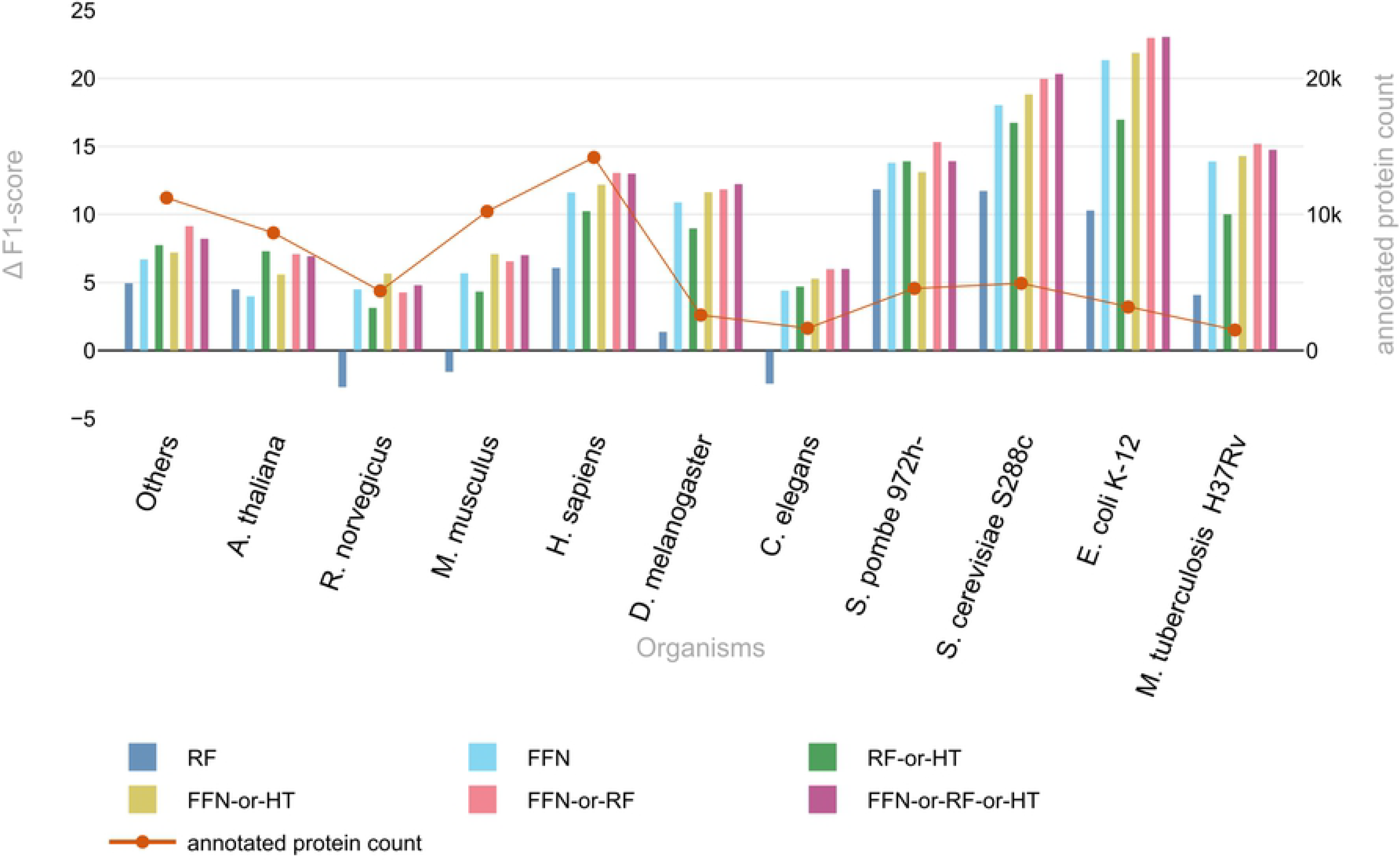
System performance on model organisms. The performance of the methods on the 10 organisms that have more than 1,000 annotated protein sequences in the training data. *Others* represents a group of organisms with less than 1,000 annotated protein sequences. The vertical bars plot the difference of each tested method to the Homology Transfer fallback (left vertical scale). The red connecting line plots the number of annotated proteins for each organism (right vertical scale).

Considering the different ontologies, predicting biological processes remains a challenge for the methods, compared to cellular component and molecular function. As shown in Fig 3, all of the ensembles exhibit the highest performance when predicting cellular components, followed by molecular function and biological process, a trend common to many AFP methods [2, 13, 18]. The difficulty of predicting *biological process* terms is probably due to the fact that its terms are the least correlated with sequence similarity [48, 49]. Thus using only sequence similarity to infer functions can be insufficient or misleading, e.g. paralogs which occur from evolutionary gene duplication processes are often recruited to different pathways [50].

**Fig 3.**
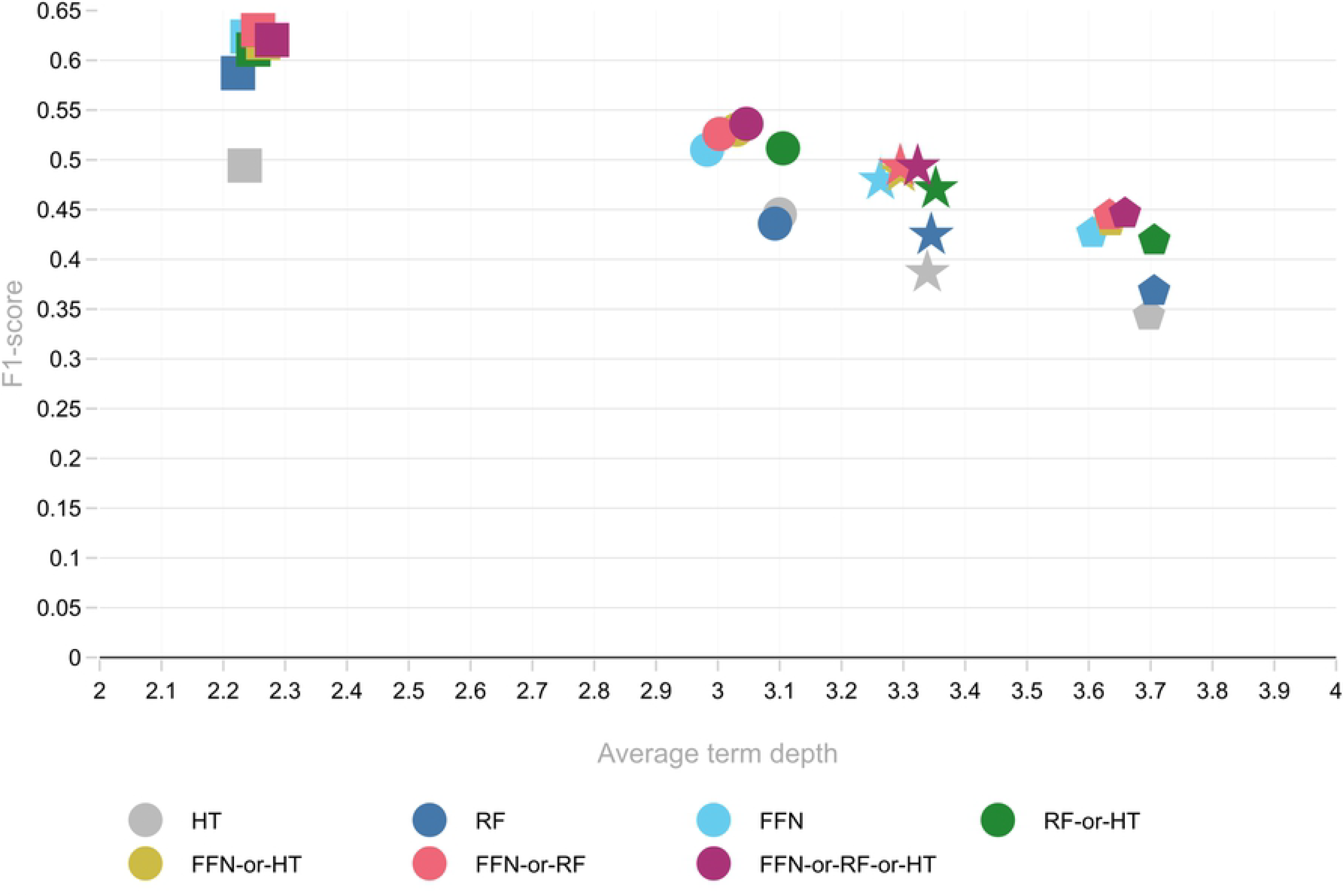
System performance on each ontology. The performance of the systems for each model on cellular component (square), molecular function (circle), biological process (pentagon) and all ontologies combined (star). The average term depths are calculated from the predictions in each ontology.

Fig 3 also shows a near-linear dependence of the F-score on the average term depth in the three GO ontologies, which correlates with the complexity and richness of these ontologies. Overall, the FFN predictions seem marginally less specific as the average term depth of FFN predictions is lower by 0.1 compared to the HT and RF approaches.

In the official evaluation of the CAFA3 challenge our ensemble system has demonstrated good overall performance, ranking among the top-10 methods for all three ontologies, with a particularly strong performance for molecular functions and cellular component of prokaryotic proteins (top-3). At the time of writing, only the top-10 systems can be compared, since detailed numerical results for all teams are not available.

## Conclusion

We have presented methods for automated protein function prediction, evaluated both on Swiss-Prot and also through the CAFA3 challenge. These methods have demonstrated competitive performance among more than 100 CAFA3 entries, especially on the prediction of prokaryotic molecular functions and cellular components. Nevertheless, the absolute performance of the AFP methods show that the task remains a challenge.

We can chart two main directions for further development. Firstly, features derived from sequences related in other ways than solely through homology, e.g. through co-expression or binding, can be potentially beneficial especially for the prediction of biological processes, as demonstrated for instance by Piovesan et al. [51] and Kulmanov et al. [18]. Of the three GO ontologies, biological process currently exhibits the lowest absolute performance, and therefore is the most impactful target for further development.

Secondly, structure and sequence-based features without doubt play an important role in determining the function of a protein. However, having to employ the large number of external tools needed to obtain the relevant features is a surprisingly tedious task. As a potential remedy to this practical problem, but also as a research task in its own right, we experimented with using a CNN to derive features directly from the sequence, without any external analysis tools. While the absolute performance of the CNN method can not currently compete with the feature-based methods, the CNN achieved what we believe to be a surprisingly good performance given the simple format of its input. The CNN performing on par with purely BLAST-based predictions suggests that, with further development, the reliance on homology — an important source of features in much of the current AFP work — could potentially be omitted, with a neural model analyzing the amino acid sequences directly. However, training a well-performing neural model is non-trivial. As future work, we plan to improve our methods by testing other neural network architectures [18, 21] and by pretraining the used protein sequence encoders in a similar fashion as is common in neural computer vision and natural language processing systems [52, 53]

The trained prediction models and source code of the system are publicly available under an open license at https://github.com/TurkuNLP/CAFA3.

## Supporting information

**S1 Table. Feature group selection.** Feature groups are selected by removing one more at each round (RM). Performance is shown as micro-averaged F-score (F), precision (P) and recall (R), evaluated for the top-5000 terms. After removing three groups performance no longer increases. Performance is measured on the *optimization* set. Performance on the *test* set also increases from 0.411 for not being over-fitted on the *optimization* set.

**S2 Table. Ensemble system performance.** The ensemble system is made by combining the predictions from the two machine learning systems (RF = random forest classifier and FFN = feedforward neural network) and the homology transfer (HT) with either boolean *AND* or boolean *OR*. Performance is shown as micro-averaged F-score (F), precision (P) and recall (R). F ∆ shows the difference to the best performing system combination measured in percentage points.

## Acknowledgments

We would like to thank Sampo Pyysalo for his insightful comments on the manuscript. We also thank CSC - IT Center for Science Ltd, Espoo, Finland for providing the computational resources used in this work.

## References

1. Radivojac P, Clark WT, Oron TR, Schnoes AM, Wittkop T, Sokolov A, et al. A large-scale evaluation of computational protein function prediction. Nature methods. 2013;10(3):221.

2. Jiang Y, Oron TR, Clark WT, Bankapur AR, D’Andrea D, Lepore R, et al. An expanded evaluation of protein function prediction methods shows an improvement in accuracy. Genome biology. 2016;17(1):184.

3. The Gene Ontology Consortium. The Gene Ontology Resource: 20 years and still GOing strong. Nucleic Acids Research. 2018;47(D1):D330–D338. doi:10.1093/nar/gky1055.

4. Ashburner M, Ball CA, Blake JA, Botstein D, Butler H, Cherry JM, et al. Gene ontology: tool for the unification of biology. Nature genetics. 2000;25(1):25.

5. The UniProt Consortium. UniProt: the universal protein knowledgebase. Nucleic Acids Research. 2017;45(D1):D158–D169. doi:10.1093/nar/gkw1099.

6. Cozzetto D, Buchan DW, Bryson K, Jones DT. Protein function prediction by massive integration of evolutionary analyses and multiple data sources. BMC Bioinformatics. 2013;14(3):S1. doi:10.1186/1471-2105-14-S3-S1.

7. Lipman DJ, Zhang J, Madden TL, Altschul SF, Schäffer AA, Miller W, et al. Gapped BLAST and PSI-BLAST: a new generation of protein database search programs. Nucleic Acids Research. 1997;25(17):3389–3402. doi:10.1093/nar/25.17.3389.

8. Nokso-Koivisto J, Holm L, Koskinen P, Törönen P. PANNZER: high-throughput functional annotation of uncharacterized proteins in an error-prone environment. Bioinformatics. 2015;31(10):1544–1552. doi:10.1093/bioinformatics/btu851.

9. Altschul SF, Gish W, Miller W, Myers EW, Lipman DJ. Basic local alignment search tool. Journal of Molecular Biology. 1990;215(3):403–410. doi:https://doi.org/10.1016/S0022-2836(05)80360-2.

10. Falda M, Toppo S, Pescarolo A, Lavezzo E, Di Camillo B, Facchinetti A, et al. Argot2: a large scale function prediction tool relying on semantic similarity of weighted Gene Ontology terms. BMC Bioinformatics. 2012;13(4):S14. doi:10.1186/1471-2105-13-S4-S14.

11. Eddy SR. Profile hidden Markov models. Bioinformatics. 1998;14(9):755–763. doi:10.1093/bioinformatics/14.9.755.

12. Gong Q, Ning W, Tian W. GoFDR: A sequence alignment based method for predicting protein functions. Methods. 2016;93:3–14. doi:https://doi.org/10.1016/j.ymeth.2015.08.009.

13. You R, Zhang Z, Xiong Y, Sun F, Mamitsuka H, Zhu S. GOLabeler: improving sequence-based large-scale protein function prediction by learning to rank. Bioinformatics. 2018;34(14):2465–2473. doi:10.1093/bioinformatics/bty130.

14. Mitchell AL, Sangrador-Vegas A, Luciani A, Madeira F, Nuka G, Salazar GA, et al. InterPro in 2019: improving coverage, classification and access to protein sequence annotations. Nucleic Acids Research. 2018;47(D1):D351–D360. doi:10.1093/nar/gky1100.

15. Ofer D, Linial M. ProFET: Feature engineering captures high-level protein functions. Bioinformatics. 2015;31(21):3429–3436. doi:10.1093/bioinformatics/btv345.

16. Hou J, Adhikari B, Cheng J. DeepSF: deep convolutional neural network for mapping protein sequences to folds. Bioinformatics. 2017;34(8):1295–1303.

17. Wang J, Cao H, Zhang JZ, Qi Y. Computational protein design with deep learning neural networks. Scientific reports. 2018;8(1):6349.

18. Kulmanov M, Khan MA, Hoehndorf R. DeepGO: predicting protein functions from sequence and interactions using a deep ontology-aware classifier. Bioinformatics. 2017;34(4):660–668. doi:10.1093/bioinformatics/btx624.

19. Roth A, Franceschini A, Szklarczyk D, Heller D, Simonovic M, Wyder S, et al. STRING v10: protein-protein interaction networks, integrated over the tree of life. Nucleic Acids Research. 2014;43(D1):D447–D452. doi:10.1093/nar/gku1003.

20. Cao R, Freitas C, Chan L, Sun M, Jiang H, Chen Z. ProLanGO: protein function prediction using neural machine translation based on a recurrent neural network. Molecules. 2017;22(10):1732.

21. Fa R, Cozzetto D, Wan C, Jones DT. Predicting human protein function with multi-task deep neural networks. PLOS ONE. 2018;13(6):1–16. doi:10.1371/journal.pone.0198216.

22. Teng S, Luo H, Wang L. Random forest-based prediction of protein sumoylation sites from sequence features. In: Proceedings of the First ACM International Conference on Bioinformatics and Computational Biology. ACM; 2010. p. 120–126.

23. Jo T, Cheng J. Improving protein fold recognition by random forest. In: BMC bioinformatics. vol. 15. BioMed Central; 2014. p. S14.

24. Jain P, Hirst JD. Automatic structure classification of small proteins using random forest. BMC Bioinformatics. 2010;11(1):364. doi:10.1186/1471-2105-11-364.

25. Liu M, Chen XW. Prediction of protein-protein interactions using random decision forest framework. Bioinformatics. 2005;21(24):4394–4400. doi:10.1093/bioinformatics/bti721.

26. Kumar C, Li G, Choudhary A. Enzyme function classification using protein sequence features and random forest. In: 2009 3rd International Conference on Bioinformatics and Biomedical Engineering. IEEE; 2009. p. 1–4.

27. Zhou N, Jiang Y, Bergquist TR, Lee AJ, Kacsoh BZ, Crocker AW, et al. The CAFA challenge reports improved protein function prediction and new functional annotations for hundreds of genes through experimental screens. bioRxiv. 2019;doi:10.1101/653105.

28. Camacho C, Coulouris G, Avagyan V, Ma N, Papadopoulos J, Bealer K, et al. BLAST+: architecture and applications. BMC bioinformatics. 2009;10(1):421.

29. Henikoff S, Henikoff JG. Amino acid substitution matrices from protein blocks. Proceedings of the National Academy of Sciences. 1992;89(22):10915–10919. doi:10.1073/pnas.89.22.10915.

30. Boratyn GM, Schäffer AA, Agarwala R, Altschul SF, Lipman DJ, Madden TL. Domain enhanced lookup time accelerated BLAST. Biology Direct. 2012;7(1):12. doi:10.1186/1745-6150-7-12.

31. Zheng C, Lanczycki CJ, Zhang D, Hurwitz DI, Chitsaz F, Lu F, et al. CDD/SPARCLE: functional classification of proteins via subfamily domain architectures. Nucleic Acids Research. 2016;45(D1):D200–D203. doi:10.1093/nar/gkw1129.

32. Jones P, Binns D, Chang HY, Fraser M, Li W, McAnulla C, et al. InterProScan 5: genome-scale protein function classification. Bioinformatics. 2014;30(9):1236–1240.

33. Federhen S. The NCBI taxonomy database. Nucleic acids research. 2011;40(D1):D136–D143.

34. Heddad A, Brameier M, MacCallum RM. Evolving regular expression-based sequence classifiers for protein nuclear localisation. In: Workshops on Applications of Evolutionary Computation. Springer; 2004. p. 31–40.

35. Kiemer L, Bendtsen JD, Blom N. NetAcet: prediction of N-terminal acetylation sites. Bioinformatics. 2005;21(7):1269–1270. doi:10.1093/bioinformatics/bti130.

36. Pierleoni A, Martelli PL, Casadio R. PredGPI: a GPI-anchor predictor. BMC bioinformatics. 2008;9(1):392.

37. Tung CH, Chen CW, Sun HH, Chu YW. Predicting human protein subcellular localization by heterogeneous and comprehensive approaches. PLOS ONE. 2017;12(6):1–14. doi:10.1371/journal.pone.0178832.

38. Kawashima S, Pokarowski P, Pokarowska M, Kolinski A, Katayama T, Kanehisa M. AAindex: amino acid index database, progress report 2008. Nucleic Acids Research. 2008;36(suppl 1):D202–D205. doi:10.1093/nar/gkm998.

39. Srivastava N, Hinton G, Krizhevsky A, Sutskever I, Salakhutdinov R. Dropout: a simple way to prevent neural networks from overfitting. The Journal of Machine Learning Research. 2014;15(1):1929–1958.

40. Breiman L. Random forests. Machine learning. 2001;45(1):5–32.

41. Pedregosa F, Varoquaux G, Gramfort A, Michel V, Thirion B, Grisel O, et al. Scikit-learn: Machine Learning in Python. Journal of Machine Learning Research. 2011;12:2825–2830.

42. Krizhevsky A, Sutskever I, Hinton GE. ImageNet classification with deep convolutional neural networks. In: Advances in neural information processing systems; 2012. p. 1097–1105.

43. Zhang X, Zhao J, LeCun Y. Character-level convolutional networks for text classification. In: Advances in neural information processing systems; 2015. p. 649–657.

44. Wang S, Peng J, Ma J, Xu J. Protein secondary structure prediction using deep convolutional neural fields. Scientific reports. 2016;6:18962.

45. Xu D, Nussinov R. Favorable domain size in proteins. Folding and Design. 1998;3(1):11–17.

46. Zeng H, Edwards MD, Liu G, Gifford DK. Convolutional neural network architectures for predicting DNA–protein binding. Bioinformatics. 2016;32(12):i121–i127.

47. Blum AL, Langley P. Selection of relevant features and examples in machine learning. Artificial Intelligence. 1997;97(1):245–271. doi:https://doi.org/10.1016/S0004-3702(97)00063-5.

48. Lord PW, Stevens RD, Brass A, Goble CA. Semantic similarity measures as tools for exploring the gene ontology. In: Biocomputing 2003. World Scientific; 2002. p. 601–612.

49. Lord PW, Stevens RD, Brass A, Goble CA. Investigating semantic similarity measures across the Gene Ontology: the relationship between sequence and annotation. Bioinformatics. 2003;19(10):1275–1283.

50. Gerlt JA, Babbitt PC. Divergent Evolution of Enzymatic Function: Mechanistically Diverse Superfamilies and Functionally Distinct Suprafamilies. Annual Review of Biochemistry. 2001;70(1):209–246. doi:10.1146/annurev.biochem.70.1.209.

51. Piovesan D, Giollo M, Ferrari C, Tosatto SCE. Protein function prediction using guilty by association from interaction networks. Amino Acids. 2015;47(12):2583–2592. doi:10.1007/s00726-015-2049-3.

52. Kornblith S, Shlens J, Le QV. Do better ImageNet models transfer better? arXiv preprint arXiv:180508974. 2018;.

53. Devlin J, Chang MW, Lee K, Toutanova K. BERT: Pre-training of deep bidirectional transformers for language understanding. arXiv preprint arXiv:181004805. 2018;.

